# New self-identities evolve via point mutation in an invertebrate allorecognition gene

**DOI:** 10.1101/2020.12.01.406256

**Authors:** Aidan Huene, Traci Chen, Matthew L. Nicotra

## Abstract

Many organisms use genetic self-recognition systems to distinguish themselves from other members of their species. To understand how new self-identities evolve, we studied Allorecognition 2 (Alr2), a self-recognition gene from the colonial cnidarian, *Hydractinia symbiolongicarpus*. *Alr2* encodes a highly polymorphic transmembrane protein that discriminates self from non-self by selectively binding across cell membranes to other Alr2 proteins with identical or very similar sequences. Here, we show that new Alr2 proteins evolve by amino acid substitutions that immediately create isoforms with entirely novel binding specificities, or through intermediates with relaxed binding specificities. Our results also suggest a topology for homophilic interactions between Alr2 proteins. These results provide direct evidence for the generation and maintenance of functional variation at an allorecognition locus and reveal that one-component and two-component self-recognition systems evolve via different mechanisms.

## Introduction

The ability to discriminate self from same-species non-self has evolved in plants (Fujii et al., 2016), fungi (Gonçalves et al., 2020; Paoletti, 2016), slime molds (Kundert and Shaulsky, 2019), marine invertebrates (Nicotra, 2019), and bacteria (Cao et al., 2019; Gibbs and Greenberg, 2011; Pathak et al., 2013). In all cases, this ability is based on an organism’s genotype at polymorphic self-recognition loci. This polymorphism is thought to be maintained by a form of balancing selection called negative frequency-dependent selection (Kimura and Crow, 1964; Wright, 1939). Under negative frequency-dependent selection, alleles become more fit as they become less frequent. This is because rare alleles are unlikely to be shared by chance, making them better markers of self. New alleles, the rarest of all, spread in a population until their frequencies reach that of other alleles (Richman and Kohn, 2000). These dynamics can maintain tens to hundreds of self-recognition alleles in a population (e.g. Casselton and Olesnicky, 1998; Gloria-Soria et al., 2012; Goncalves et al., 2019; James, 2015; Lawrence, 2000; Nydam et al., 2017). How so many self-recognition alleles can evolve remains an evolutionary puzzle.

Most work on the evolution of new self-recognition alleles has focused on multi-component systems such as plant self-incompatibility loci, in which discrimination is controlled by linked genes encoding proteins that have co-evolved to interact (or not interact) with each other (Bod’ova et al., 2018; Chantreau et al., 2019; Chookajorn et al., 2004; Matton et al., 1999; Uyenoyama et al., 2001). In contrast, the evolution of new alleles in one-component systems has received little attention. In one-component systems, self-recognition proteins act as their own ligands. Each allele thus encodes a protein that interacts with itself but not those encoded by alternative alleles. This requirement places different constraints on the evolution of new alleles, and may even make it less likely for one-component self-recognition systems to evolve (De Tomaso, 2006).

The self-recognition gene *Allorecognition 2* (*Alr2*) from the colonial cnidarian *Hydractinia symbiolongicarpus* is an example of a one-component self-recognition system. *Alr2* encodes a transmembrane protein that binds to itself across opposing cell membranes (Karadge et al., 2015; Nicotra et al., 2009). Binding is restricted to allelic isoforms with identical or very similar sequences, and can be prevented by amino acid differences in its N-terminal Ig-like domain (domain 1) (Karadge et al., 2015). This domain is highly polymorphic, with 183 distinct amino acid sequences being maintained in a single population (Gloria-Soria et al., 2012). These results, combined with the fact that *Hydractinia* must have matching *Alr2* alleles to recognize each other as self, have led to the hypothesis that homophilic binding of Alr2 is a mechanism of self-recognition *in vivo*.

Here, we sought to understand how new *Alr2* alleles evolve. To do so, we identified a clade of five domain 1 isoforms that differed by six or fewer amino acids. We then used ancestral sequence reconstruction and *in vitr*o binding assays to determine how their specificities changed over time. Our results demonstrate that alleles with new specificities can evolve via single amino acid changes or through intermediates with broadened specificities. Finally, we combine these results with predicted three-dimensional structures and population level polymorphism data to suggest Alr2 protein-protein interactions occur in a side-to-side manner.

## Results

### Point mutations in domain 1 can create new binding specificities

We searched a dataset of full-length, naturally occurring *Alr2* alleles (Gloria-Soria et al., 2012; Nicotra et al., 2009), and identified two (111A06 and 214E06) that encoded Alr2 allelic isoforms (hereafter, “isoforms”) with six amino acid differences in domain 1 and identical sequences across the rest of the extracellular region (Fig 1A, B). Using cell aggregation assays (Karadge et al., 2015), we found that each isoform bound to itself across opposing cell membranes but did not bind to the other (Fig 1C). We therefore sought to identify the amino acid differences that prevented them from binding to each other.

**Figure 1.**
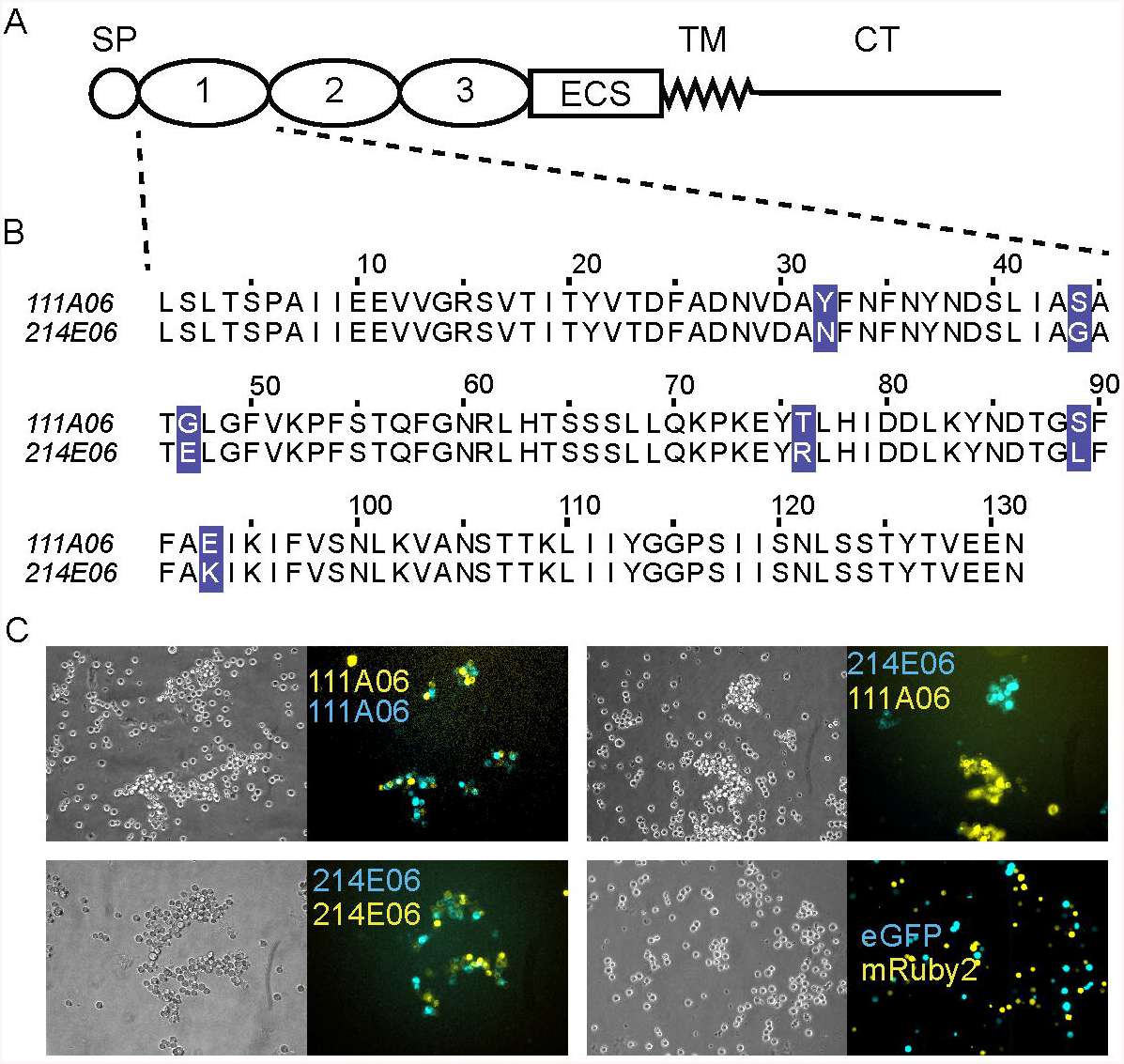
Isoform-specific, homophilic binding of Alr2 isoforms. A) Alr2 protein structure. SP = Signal peptide, ECS = Extracellular spacer, TM = Transmembrane domain, CT = Cytoplasmic tail. B) Multiple sequence alignment of 111A06 and 214E06 domain 1. Polymorphisms highlighted in purple. C) Cell aggregation assays of 111A06 and 214E06. Cells transfected with vectors encoding only fluorescent proteins (eGFP or mRuby2) do not form aggregates (bottom right).

Each amino acid difference between 111A06 and 214E06 is the result of one point mutation. To reconstruct the evolutionary history of these mutations, we created a phylogeny of all known domain 1 coding sequences (Fig 2A). 111A06 and 214E06 were located in a clade with three additional sequences (Fig 2B). We then used ancestral sequence reconstruction to infer the sequence of each node. All but the ancestral node (Anc) were predicted to be identical to an extant sequence (Fig 2C). Because 214E06 and Hap010 differed only by a single synonymous mutation, we used 214E06 to represent their shared amino acid sequence.

**Figure 2.**
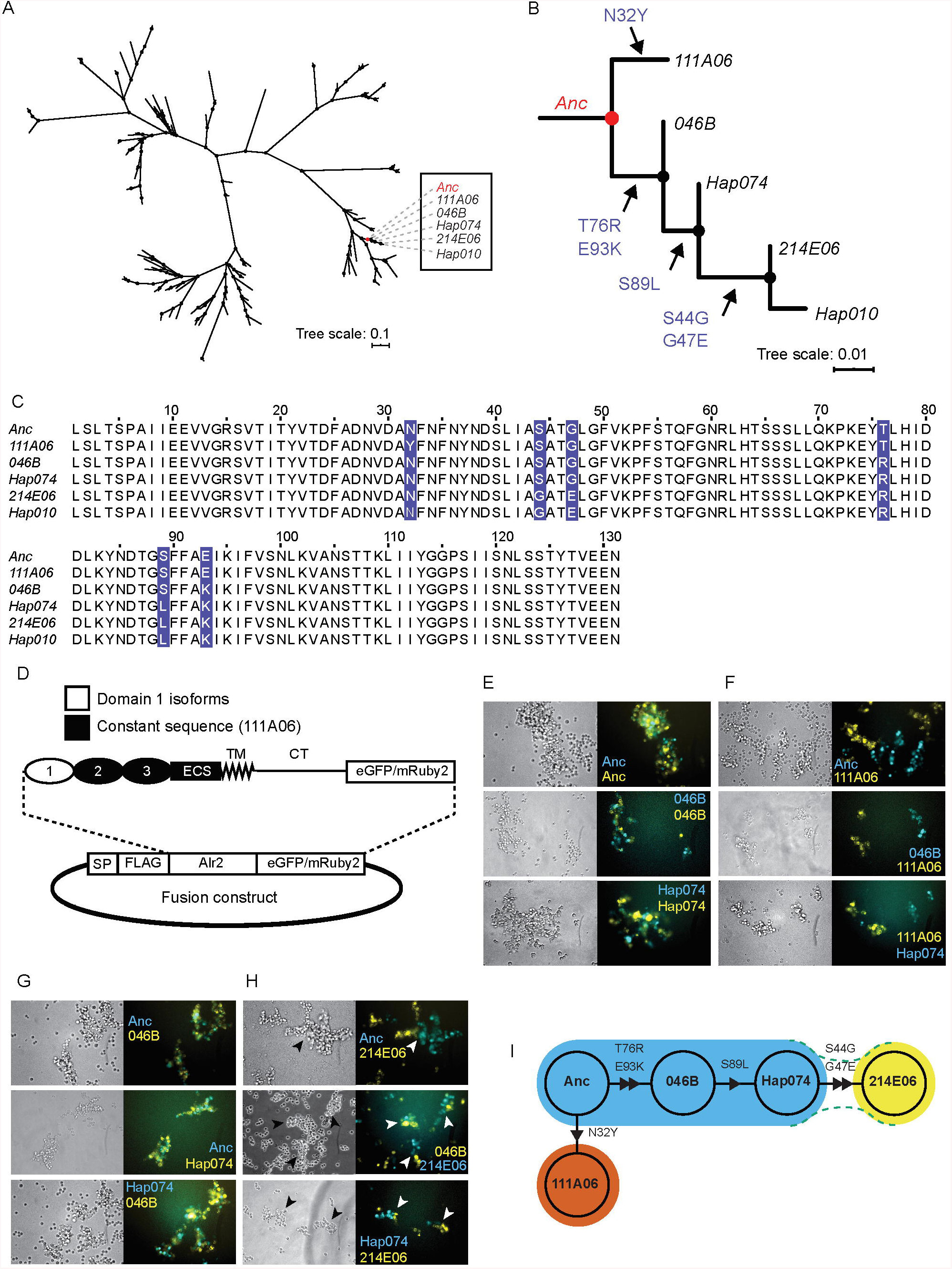
Evolution of novel binding specificities via point mutation. A) Maximum-likelihood tree of 146 domain 1 coding sequences. B) Expansion of clade that includes 111A06 and 214E06. Allele names on branch. Amino acid changes indicated along branches. C) Multiple sequence alignment of clade. Variant residues highlighted. D) Plasmid map Alr2 fusion proteins. E-H) Representative images of cell aggregation assays. E) Anc, 046B, and Hap074 against themselves. F) Anc, 046B, and Hap074 against 111A06. G) All pairwise combinations of Anc, 046B, and Hap074. H) Anc, 046B, and Hap074 versus 214E06. Arrowheads point to semi-mixed aggregates (See also Fig S1). I) Node network of isoforms colored by binding specificity. Triangles indicate the hypothesized direction of mutation from Anc. Green dotted lines indicate weaker heterophilic interactions.

To determine the binding specificity of the domain 1 isoforms encoded by these sequences, we expressed each as a fusion to domain 2 through the cytoplasmic tail of the 111A06 isoform, with a C-terminal fluorescent protein tag (Fig 2D). The resulting isoforms were tested against themselves and each other in cell aggregation assays (Fig 2E-H). Each isoform, including the predicted ancestor, Anc, caused cells to form multicellular aggregates, indicating it was capable of homophilic binding (Fig 2E). In pairwise assays, 111A06 did not form mixed aggregates with any isoform, indicating it had a unique binding specificity within the clade (Fig 2F). In contrast, Anc, 046B, and Hap074 all formed mixed aggregates with each other, indicating a shared binding specificity (Fig 2G).

In assays that paired 214E06 with Anc, 046B, or Hap074, we observed single-color aggregates, some of which appeared to adhere to aggregates of a different color (Fig 2H, arrowheads). These semi-mixed aggregates were repeatable (Fig S1A) and qualitatively different from the mixed aggregates it formed when paired with itself, and from the completely separate aggregates it formed with the other four isoforms. This ruled out a defect in 214E06 that prevented homophilic binding or caused it to bind to any isoform. Semi-mixed aggregates have been observed in studies of cell adhesion molecules that have strong homophilic affinities, but weaker heterophilic affinities (Goodman et al., 2016; Katsamba et al., 2009). Because of this, we concluded that 214E06 binds more weakly to Anc, 046B, and Hap074 than to itself, and that it therefore had a different binding profile from the other isoforms.

Our results are consistent with the following evolutionary history (Figure 2I). An ancestral sequence, Anc, underwent a single mutation, N32Y, which created a daughter sequence, 111A06, with a novel binding specificity. In a separate lineage, the Anc sequence underwent two mutations, T76R and E93K to create 046B, which retained the ability to bind to Anc. A third mutation, S89L, then created Hap074, which also remained able to bind Anc and 046B. Two more mutations, S44G and G47E, then created 214E06, which bound more weakly to the ancestral isoforms than to itself (Fig 2I, dotted lines). The result is a clade in which we can discern three binding specificities, one of which arose via a single point mutation.

### New alleles can evolve via intermediates with broad specificities

Within the phylogeny, two pairs of mutations occurred within single branches (Fig 2B), preventing us from determining which came first. To determine whether the missing single-step intermediates were functional (i.e., able to bind homophilically) or had a different binding specificity from their parent and daughter sequences, we re-created each one (Fig 3A) and tested it in cell aggregation assays. We found each intermediate could bind homophilically (Fig 3B), thus ruling out the possibility that there were non-functional intermediates in the clade.

**Figure 3.**
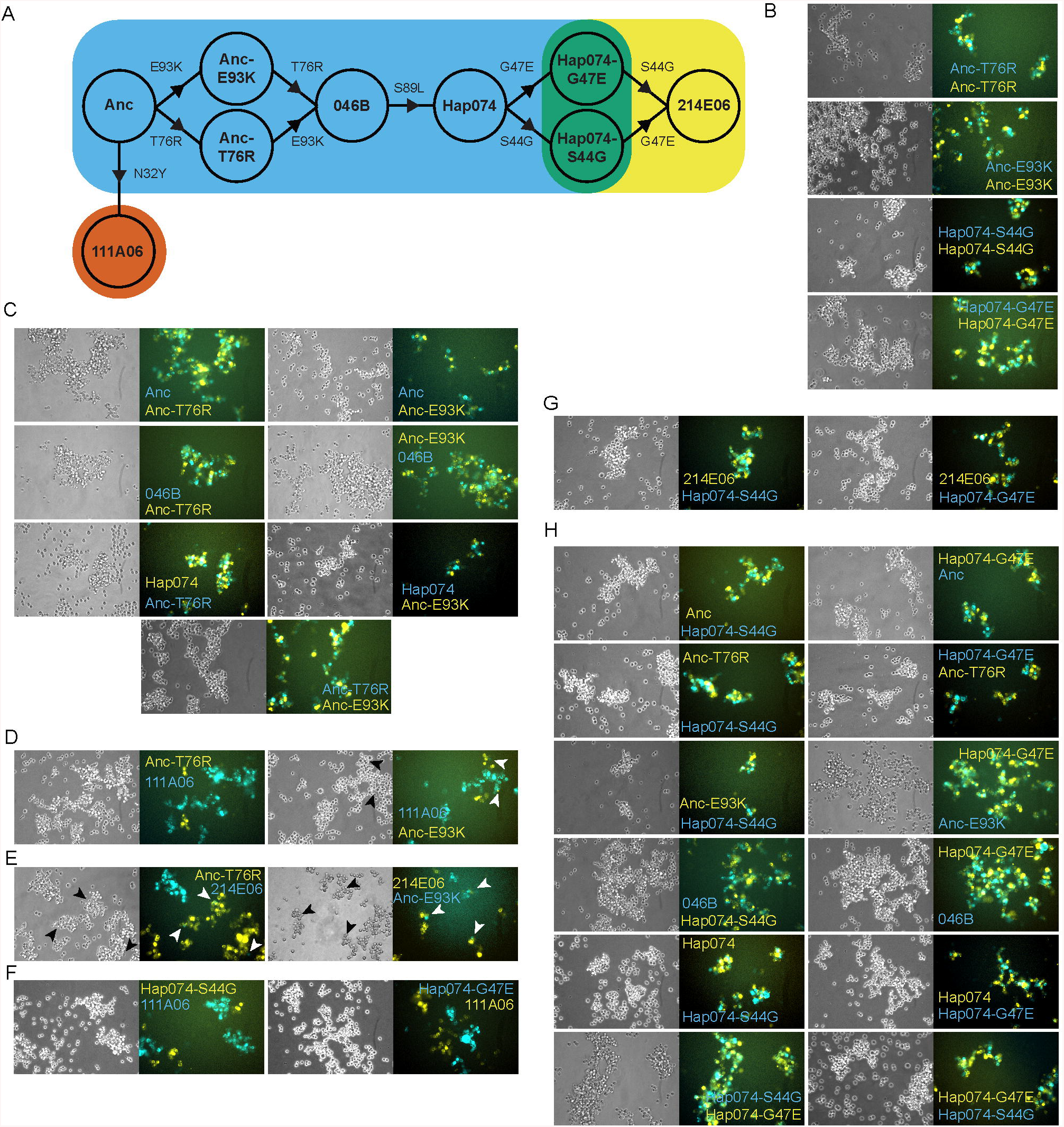
Domain 1 isoforms can evolve via intermediates with broadened specificity. A) Expanded node network including hypothesized single-step mutants between Anc and 046B, Hap074 and 214E06. B-H) Representative images of cell aggregation assays. B) Mutants tested against themselves. C) Anc-T76R and Anc-E93K tested against Anc, 046B, Hap074. D) Anc-T76R versus 111A06 (left) and Anc-E93K versus 111A06 (right). Semi-mixed aggregates indicated with arrowheads (See also Fig S1A). E) Anc-T76R and Anc-E93K versus 214E06 (See also Fig S1C). F) Hap074-S44G and Hap074-G47E versus 111A06. G) Hap074-S44G and Hap074-G47E versus 214E06. H) Hap074-S44G and Hap074-G47E versus all remaining isoforms.

We next tested the specificity of each missing intermediate. The first pair, Anc-T76R and Anc-E93K, formed mixed aggregates with Anc, 046B, and Hap074 (Fig 3C). Assays pairing Anc-T76R with 111A06 resulted in single color aggregates (Fig 3D), but those pairing Anc-E93K with 111A06 resulted in a few semi-mixed aggregates (Fig 3D, arrowheads, Fig S1B). Both mutants also formed semi-mixed aggregates when paired with 214E06 (Fig 3E, Fig S1C). Thus, evolution from Anc to 046B is unlikely to have involved a significant change in binding specificity (Fig 3A).

In contrast, the specificity of the second pair of intermediates, Hap074-S44G and Hap074-G47E, was different from their parent and daughter sequences. These mutants failed to form mixed aggregates with 111A06 (Fig 3F) but did form mixed aggregates with 214E06 (Fig 3G) and all other ancestral sequences (Fig 3H). We did not observe semi-mixed aggregates in any assay. These results suggest the first mutation on the path from Hap074 to 214E06 created a sequence that could still bind Hap074 (Fig 3A). A second mutation then generated a new allele, 214E06, which remained able to bind its parent sequence, but had a weaker affinity for Hap074. The evolution of new domain 1 sequences can therefore proceed through intermediates with broader specificities than their parental or daughter sequences.

### The N32Y mutation preserves homophilic binding and alters specificity

Isoform 111A06 evolved when position 32 mutated from Asp to Tyr in Anc. We therefore hypothesized the N32Y mutation might turn 046B or Hap074, which had the same specificity as Anc, into isoforms with the same specificity as 111A06. To test this, we generated 046B-N32Y and Hap074-N32Y. In assays with themselves, each formed mixed aggregates, indicating the mutation did not disrupt homophilic binding (Fig S1D). In pairwise assays with each other and 111A06, the mutants formed mixed aggregates, indicating they had gained the ability to bind 111A06 and each other (Fig 4A and S1G). In pairwise assays with their immediate ancestors, however, the mutants formed semi-mixed aggregates (Fig 4B, asterisks, and Fig S1E). This indicated each could still bind its ancestor, albeit more weakly than it did itself. Finally, we performed pairwise assays with the remaining isoforms in the clade. This showed the mutants had different specificities than 111A06, 046B, or Hap074 (Fig 4B and S2). In sum, the N32Y altered the specificities of 046B and Hap074 but did not generate daughter sequences with the same specificity as 111A06.

**Figure 4.**
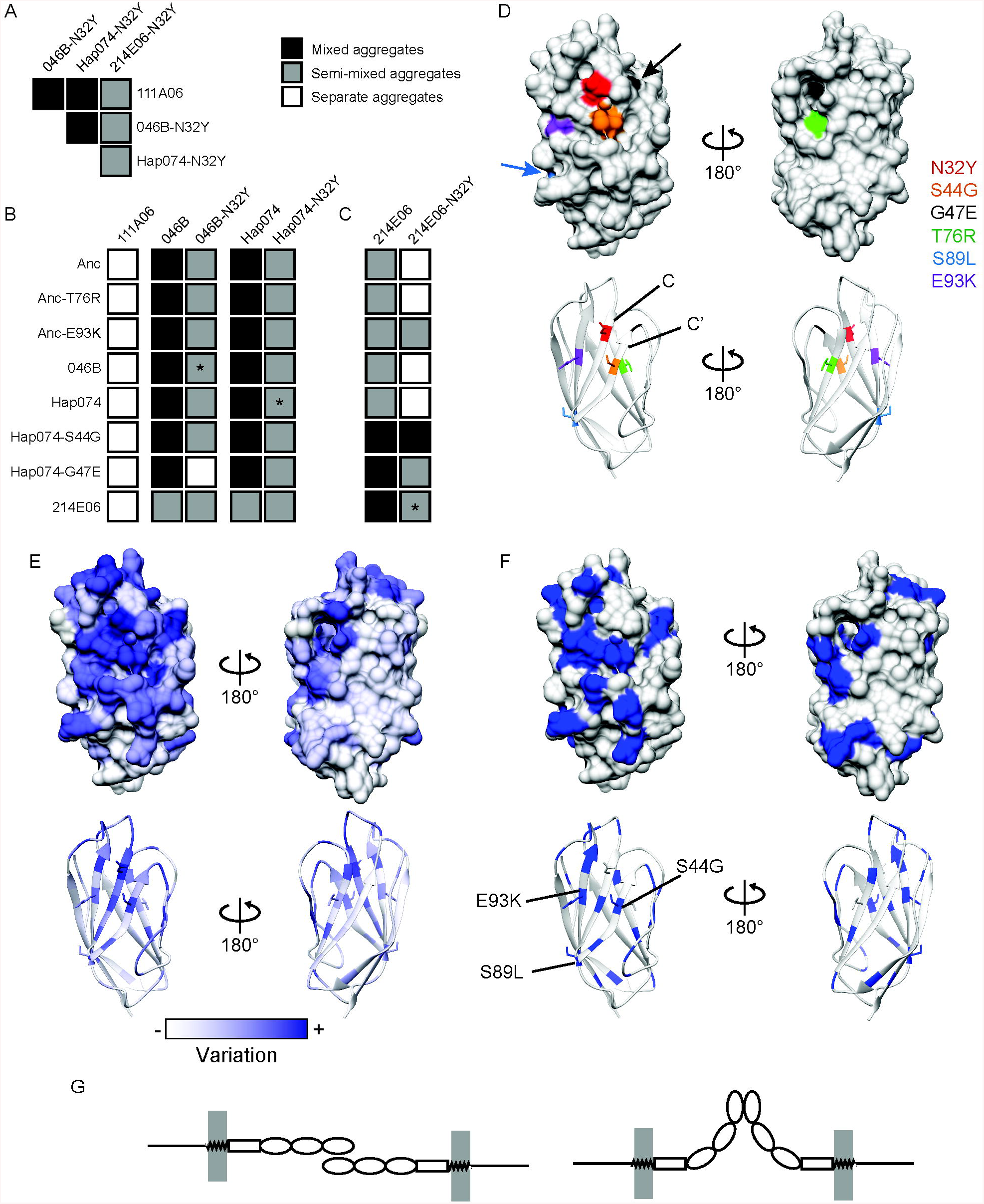
Effects of N32Y mutation on binding specificity and structural analysis. A) Results of assays between N32Y mutants and 111A06 (See also Fig S1G,H). B) Binding profiles of 111A06, 046B-N32Y, Hap074-N32Y, 046B, and Hap074. C) Binding profiles of 214E06 and 214E06-N32Y. D) Predicted structure of Anc domain 1. Six variant residues labeled. E) Sequence conservation mapped onto domain 1. F) Residues with a posterior probability ≥0.9 as determined by FUBAR are highlighted. G) Hypothetical binding topologies Alr2.

We next tested whether the N32Y mutation would alter the specificity of 214E06, the remaining domain 1 isoform known to exist in nature. We generated 214E06-N32Y and found it formed mixed aggregates with itself (Fig S1D), indicating it was able to bind homophilically. It also formed semi-mixed aggregates with 214E06 (Fig 4C, asterisk, and Fig S1F), indicating a reduced binding affinity for its immediate ancestor compared to itself. However, 214E06-N32Y only formed semi-mixed aggregates with 111A06, 046B-N32Y, and Hap074-N32Y (Fig 4A). A shared Tyr at position 32 was therefore insufficient for two isoforms to bind each other as strongly as they did themselves. Pairwise assays with the remaining isoforms revealed 214E06-N32Y to have a different binding profile than 214E06, including a gain of the ability to form mixed aggregates with Hap074-S44G (Fig 4C and S3). The effect of the N32Y mutation thus depends on the sequence context in which it occurs.

### Structural analysis of domain 1 suggests potential binding interface

In this study, three mutations altered the binding specificity of domain 1 (N32Y, S44G, and G47E), while three others did not. To investigate how these mutations might affect the tertiary structure of domain 1—and thus its binding specificity—we used I-TASSER to predict the structure of each. All were predicted to fold like a V-set Ig-domain, which was consistent with previous work (Nicotra et al., 2009). Mapping the mutations onto these structures showed five of six were predicted to reside on one face of the beta-sandwich, with the three specificity-altering mutations in close proximity to each other in beta-strands C and C’ (Fig 4D shows the structure of Anc for illustration). This suggested these strands are involved in homophilic binding between compatible domain 1 isoforms.

To gain further insight into the mechanism of homophilic binding, we then compared the I-TASSER models for domains differing by a single amino acid (e.g. Anc vs 111A06). We noted many differences in the predicted orientation of the mutated residues and their nearby amino acids. However, molecular dynamics simulations indicated these orientations were unlikely to be stable (data not shown), so we did not continue to analyze our models at this level of detail.

As an alternative approach to identify functionally important parts in domain 1, we reasoned that diversifying selection should increase sequence variation at or near the binding site. We therefore calculated the level of sequence variation at each site across all known domain 1 sequences, then mapped this metric onto the predicted structure of Anc. Most variable sites were predicted to reside on one face of the domain, which included strands C and C’ and residues 32, 44, and 47 (Fig 4E).

A second signature of diversifying selection would be increased nonsynonymous mutation rates at codons for amino acids involved in binding. To test this, we used FUBAR (Murrell et al., 2013) to identify sites under positive (diversifying) selection. Sites 44, 89, and 93 were under positive selection, while sites 32, 47, and 76 were not. Twenty-six additional sites were also under positive selection, and with fourteen of these located on the more variable face of the beta sandwich (Fig 4F).

Together, these evolutionary signatures suggest that the binding surface of domain 1 is on one side of its fold. This, in turn, suggests Alr2 proteins bind to each other via “side-to-side” interactions at their N-terminal domains. We speculate these interactions could occur in either an antiparallel or parallel topology (Fig4G).

## Discussion

*Alr2* alleles encode transmembrane proteins that discriminate between themselves and other Alr2 isoforms via homophilic binding (Karadge et al., 2015). To be favored by natural selection, new alleles must simultaneously preserve their homophilic binding capacity, lose the ability to bind old alleles, and gain the ability to bind themselves. Here, we show how new alleles evolve within this constraint. We studied a clade of closely related sequences of domain 1, a domain in which sequence differences are sufficient to prevent Alr2 proteins from binding. We show that new binding specificities can evolve by a single mutation that creates a sequence with an entirely new specificity. They can also evolve by mutations that generate an intermediate with a broader specificity, which is then constricted by additional mutations to generate a novel specificity. The fact so little molecular evolution occurred within this clade increases our confidence that our ancestral sequence reconstructions are valid. Moreover, because the sequences in this study were drawn from a single population, our results show that natural selection maintains ancestral specificities alongside novel ones. These results therefore reveal a natural process that generates, maintains, and increases functional diversity at an invertebrate self-recognition locus.

Although we focused on point mutations, other genetic mechanisms could also create *Alr2* alleles with novel specificities. Domain 1 is encoded by exon 2 of *Alr2*, which frequently recombines between *Alr2* alleles (Gloria-Soria et al., 2012). This exon also shuffles between *Alr2* and several adjacent pseudogenes by gene conversion or unequal crossing over (Gloria-Soria et al., 2012). These events could create exons encoding chimeric domain 1 sequences with novel specificities. It is also possible that protein-protein interactions between domain 1 and the rest of the extracellular region contribute to the binding specificity of a full-length Alr2 protein. If this is the case, recombining exon 2 between *Alr2* alleles with different domain 2, 3, or ECS sequences would be another way to generate new alleles.

While cell aggregation assays are commonly used to determine whether cell adhesion molecules bind to each other (Goodman et al., 2016; Kasinrerk et al., 1999; Katsamba et al., 2009; Rubinstein et al., 2015; Schreiner and Weiner, 2010; Thu et al., 2014), two caveats are germane when extrapolating our results to nature. First, these assays are qualitative and only indicate whether two isoforms bind in a shaking cell culture dish. Alleles within a binding specificity (e.g., Anc/046B/Hap074 or 111A06/046B-N32Y/Hap074-N32Y) likely have differences in affinity we could not detect. Second, the in vitro binding affinity of an Alr2 protein probably differs from its in vivo affinity. Since we do not know how strongly an Alr2 protein must bind for it to trigger self recognition in vivo, the number of distinct self-identities encoded by this clade could differ from what we have presented here. To fully resolve this issue, we will need to ectopically express these alleles in living colonies and determine their phenotypic effect, an experimental approach now possible thanks to recent advances in *Hydractinia* functional genomics (Sanders et al., 2018).

Some have argued that self-recognition systems based on homophilic binding proteins are unlikely because it is hard to imagine how new allorecognition alleles could evolve. Instead, they have hypothesized there must be a developmental process by which organisms “tune” or “educate” their self-recognition systems to recognize inherited alleles as self (Boehm, 2006; De Tomaso, 2006). While our results demonstrate that novel binding specificities can evolve via point mutation at *Alr2*, there may still be a role for an education process in *Hydractinia*. As mentioned above, *Alr2* alleles probably encode variants with different binding affinities. Moreover, some *Hydractinia* appear to have more than two *Alr2* alleles, suggesting copy-number variation exists for this gene (Gloria-Soria et al., 2012). Colonies may therefore need to set their own internal threshold—based on the complement of *Alr2* alleles they inherit—for the amount of signaling that triggers self-recognition. This might occur in developing embryos, which do not exhibit allorecognition phenomena despite constitutive expression of *Alr2* (Nicotra et al., 2009; Poudyal et al., 2007). This type of education system would be analogous NK cell licensing or allorecognition via myeloid cells in vertebrates (Dai et al., 2017, 2020; Jonsson and Yokoyama, 2009; Zamora et al., 2017).

Our results suggest that one-component self-recognition systems can evolve differently from two-component systems. At *Alr2,* for example, a single mutation generated an allele encoding a novel binding specificity that would likely allow the animal carrying it to distinguish itself from an animal carrying the ancestral allele. In contrast, at self-incompatibility loci, which encode co-evolving pairs of interacting proteins, sequential mutations in both proteins are thought to be required for them discriminate between themselves and other alleles (Chantreau et al., 2019; Chookajorn et al., 2004). This constraint may require self-incompatibility alleles to evolve through non-functional intermediates (Bod’ova et al., 2018; Uyenoyama et al., 2001), something we did not observe here. *Alr2* alleles can also evolve via dual-specificity intermediates, a mechanism that has been proposed for the evolution of new self-incompatibility alleles (Matton et al., 1999) but criticized on population genetic grounds (Charlesworth, 2000; Uyenoyama et al., 2001).

In light of our results, *Hydractinia* would be a productive system in which to study “sequence space”—the theoretical universe of all possible peptides of a given length. Long-standing questions about how many functional variants of a protein exist in sequence space, how many of these actually appear in nature, and whether evolution is constrained in its ability to reach them remain unresolved (Podgornaia and Laub, 2015; Povolotskaya and Kondrashov, 2010; Weinreich et al., 2006). Because natural selection drives the continued evolution of new allorecognition alleles, allorecognition loci like *Alr2* are essentially natural experiments exploring sequence space.

Finally, we note that Sewall Wright is widely regarded as the first to propose negative frequency dependent selection as a driver of extreme sequence polymorphism at self-recognition loci. He was inspired to do so after a colleague told him that more than 37 self-sterility alleles had evolved in a small population of evening primroses strewn across the canyons of New Mexico (Wright, 1939). While the concept of rare allele advantage has since proven valid in many systems (Aanen et al., 2008; Bod’ova et al., 2018; Cao et al., 2019; Gloria-Soria et al., 2012; Grosberg, 1988; Turner et al., 2019), his assumption that one mutation could generate a new self-identity fell out of favor as the molecular basis of self-incompatibility systems became known (Charlesworth, 2006; Richman, 2000). Our results would seem to vindicate Wright on this small point. Had he heard of hydroids instead of flowers, he would have got it entirely right.

## Acknowledgements

Molecular graphics and analyses performed with UCSF Chimera, developed by the Resource for Biocomputing, Visualization, and Informatics at the University of California, San Francisco, with support from NIH P41-GM103311. We thank Kristina Paris for assistance and discussion with running molecular dynamics for the models. M.N. was supported by NSF grant IOS-1557339. A.H. was supported by NIH T32 AI074491

## Author Contributions

A.H. and T.C. performed the aggregation assay experiments. A.H. did the data analysis and structural visualizations. A.H. and M.N. designed the experiments and wrote the paper.

## STAR Methods

### Alr2 sequence acquisition and processing

Alr2 alleles 111A06 and 214E06 were identified from previously published *Alr2* sequences (Gloria-Soria et al., 2012). To obtain a dataset of *Alr2* domain 1 sequences, we downloaded all 373 *Hydractinia symbiolongicarpus Alr2* cDNA sequences from Genbank, aligned them with MAFFT (Katoh et al., 2005) as implemented in Jalview 2.10.5 (Waterhouse et al., 2009), then trimmed the alignment leaving only the region encoding domain 1. Duplicate sequences were then removed with ElimDupes (www.hiv.lanl.gov), to yield 146 distinct domain 1 cDNA sequences, encoding 137 distinct amino acid sequences.

### Phylogenetic Analysis and Ancestral State Reconstruction

The 146 domain 1 cDNA sequences were aligned with PRANK (Löytynoja, 2014), a codon-aware alignment program (File S1). The alignment was then used to construct a phylogenetic tree using maximum likelihood through IQ-TREE (http://iqtree.cibiv.univie.ac.at/) (Trifinopoulos et al., 2016) (File S2). From the web portal, the defaults settings were used with codon selected for the sequence type, standard/universal genetic code, ultrafast bootstrap analysis with a maximum of 1000 alignments, 0.99 minimum correlation coefficient, 1000 replicates of SH-aLRT branch test, 0.5 perturbation strength, and 100 set for the IQ-TREE stopping rule. Ancestral states were estimated using the phylogenetic tree generated from IQ-TREE and the ancestral reconstruction function within PRANK (File S3) (Dutheil and Boussau, 2008; Löytynoja, 2014). An unrooted tree was generated using iTOL v5.5.1 with one iteration of equal-daylight (Letunic and Bork, 2019).

### Constructs for ectopic expression of Alr2 alleles

The plasmid backbone used for all constructs in this study was the pFLAG-CMV-3 (Sigma, E6783). Previously, it was determined that the N-terminal FLAG tag did not have an effect on the binding capability of Alr2 (Karadge et al., 2015). The *Hydractinia Alr2* allele sequences were optimized for human expression using the Integrated DNA Technologies (IDT) Codon Optimization Tool (https://www.idtdna.com/CodonOpt). The full Alr2 sequence (domain 1 in the ectodomain through the cytoplasmic tail) for 111A06 and domain 1 sequences for Anc, 046B, Hap074, and 214E06 were ordered as gBlocks Gene Fragments from IDT. All other mutant domain sequences were ordered from Twist Bioscience as Gene Fragments. Coding sequences for fluorescent proteins were cloned from vectors encoding eGFP and mRuby2 (gift from Michael Davidson, Addgene plasmid #54614). Cloning was performed using the NEBuilder HiFi DNA Assembly (New England Biolabs, E2621S) with primers designed to amplify the vector and insert sequences with ≥20 bp overlap. The FLAG-111A06-eGFP/mRuby2 plasmids (pUP801, pUP746) were cloned first and then used as the template for cloning in the other domain 1 isoforms. Within the construct, linker sequences were used before (Leu-Ala-Ala-Ala) and after (Gly-Pro-Pro-Val-Glu-Lys) the Alr2 allele.

### Expression of *Alr2* alleles in mammalian cells

HEK293T cells (ATCC Cat# CRL-3216) were cultured in accordance with ATCC guidelines. Complete HEK culture medium was made using DMEM (Fisher Science, SH30081.01), 10% fetal bovine serum (Thermofisher Scientific, #16000044), 0.001% beta-mercaptoethanol (Fisher Scientific, 21-985-023), 100 U/mL penicillin and 100 mg/mL streptomycin (Sigma, P4333-100ML). To prepare plasmids for transfection, plasmids were transformed into chemically competent bacteria and isolated from cultures using the GeneJET Plasmid Midi-prep Kit (Thermofisher Scientific, K0481) or the PureLink™ HiPure Plasmid Maxiprep Kit (Thermofisher Scientific, K2100006). Plasmids were transiently transfected into HEK293T cells using TransIT-293 (Mirus Bio, MIR 2700) according to the manufacturer’s instructions. To summarize, on day 1, HEK293T cells were plated in a 12-well plate (Fisher Scientific, #353043) at a density of 3×10^5^/well in 1 ml of complete HEK medium to achieve approximately 60-70% confluency on Day 2. On Day 2, the transfection mixture was prepared in a total volume of 100 μl using 1 ug (X μl) of plasmid DNA (plasmid concentrations between 300ng-1000ng/μl), diluted with Optimem (97-X μl), and 3 μl of TransIT-293 reagent. While incubating the DNA:lipid complexes, the cells were washed using 500 μl of DPBS (Fisher Scientific, BW17-512F), incubated with 1 ml transfection medium (complete HEK medium without antibiotics), and replaced in the 5% CO_2_ incubator. Once the DNA:lipid complexes had incubated, the 100 μl mixture was added to the appropriate well, the plate gently shaken back and forth and then replaced in the incubator. On Day 4, cells were used in the aggregation assay.

### Aggregation assay

Our aggregation protocol is adapted from previous work (Karadge et al., 2015). To summarize, previously transfected HEK293T cells were incubated with 0.25% Trypsin/0.1% EDTA solution (Corning, MT25053CI), washed in complete HEK culture medium, mechanically disrupted via pipette, and filtered through a 35μm strainer mesh (Steller scientific, FSC-FLTCP) to create a single cell suspension. For each aggregation assay, a total of 5×10^4^ cells were resuspended in 500 μl aggregation assay medium (complete HEK medium, 70 U/ml DNase I [Sigma, D4527-10KU], and 2 mM EGTA [Goldbio, E-217-25]) and added to one well of a 24-well ultra-low attachment plate (Fisher Scientific, 07-200-602). When testing isoforms pairwise, 2.5×10^4^ cells of each transfection were added to the same well and resuspended in a total of 500 μl. The plate was incubated for one hour at 37°C in 5% CO_2_ on an orbital rotator (IBI Scientific, Model# BBUAAUVIS) set at 90 rpm. Assays were visualized using an inverted fluorescence microscope (Nikon Eclipse TS100). Each pairwise assay was repeated at least three times. In cases when the assay results could not be viewed immediately, cell aggregates were fixed by adding 500 μl of 8% paraformaldehyde (Fisher, AA433689M) diluted in DPBS to each well and the results imaged within 5 hrs.

### Sequence Variability and Visualization of Domain 1

The structure for the Anc domain 1 isoform was predicted using I-TASSER v5.1 (Roy et al., 2010; Yang et al., 2015; Zhang, 2008) which resulted in a domain with a V-set like fold. To visualize the variable positions within domain 1, the aligned 137 protein sequences were uploaded to the Multialign Viewer in UCSF Chimera (Meng et al., 2006; Pettersen et al., 2004) and the conservation rendered onto the structure. Sites under positive selection were identified using FUBAR (Murrell et al., 2013) through the HyPhy (Pond et al., 2005, 2019) and Datamonkey (http://www.datamonkey.org/) (Delport et al., 2010; Pond and Frost, 2005; Weaver et al., 2018) platform. The sites under positive selection (posterior probability of ≥0.9) were mapped onto domain 1.

**Supplemental Information**

