## Supplementary figures and images for "New self-identities evolve via point mutation in an invertebrate allorecognition gene"

### Figure S1

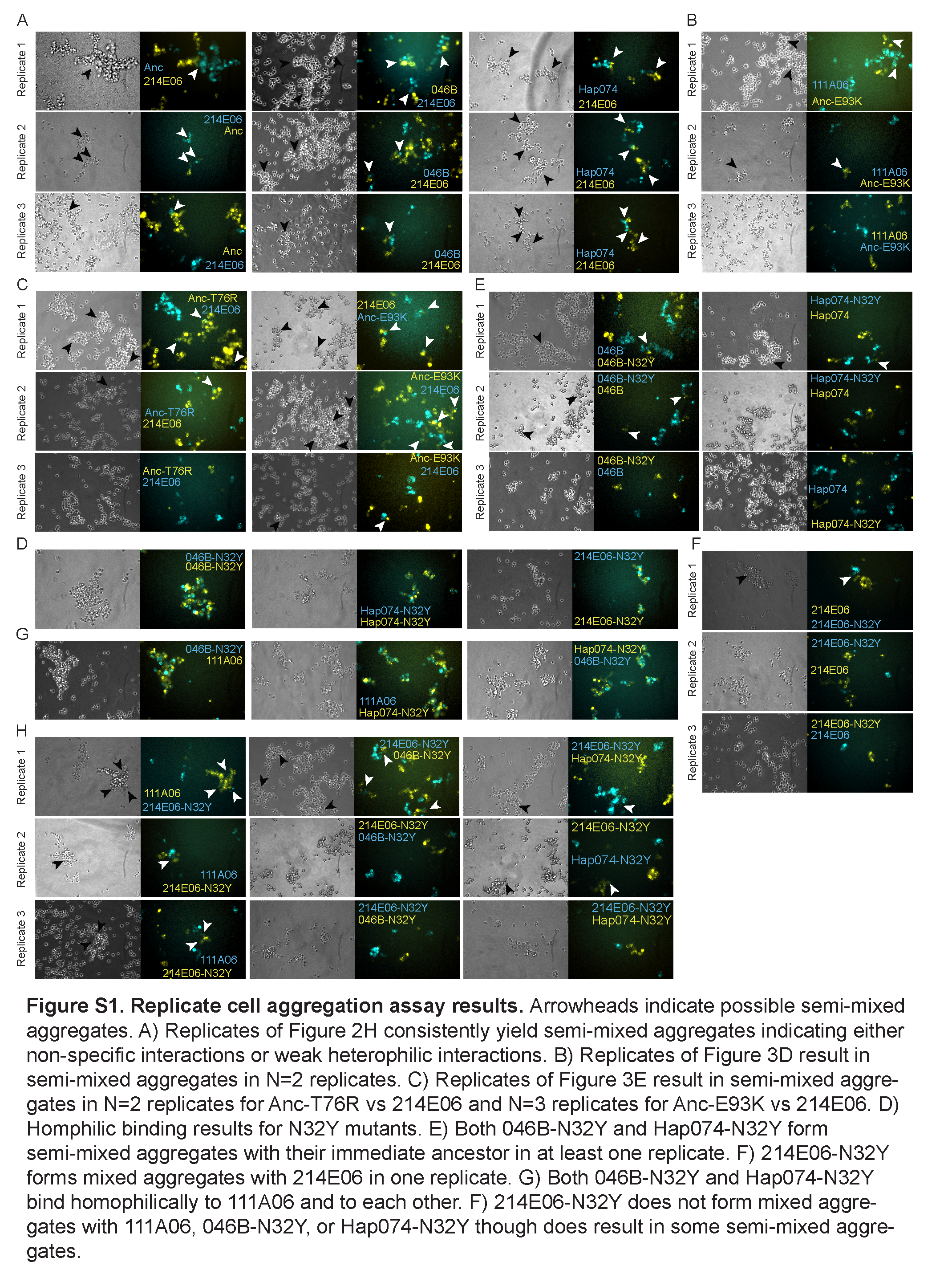

### Figure S2

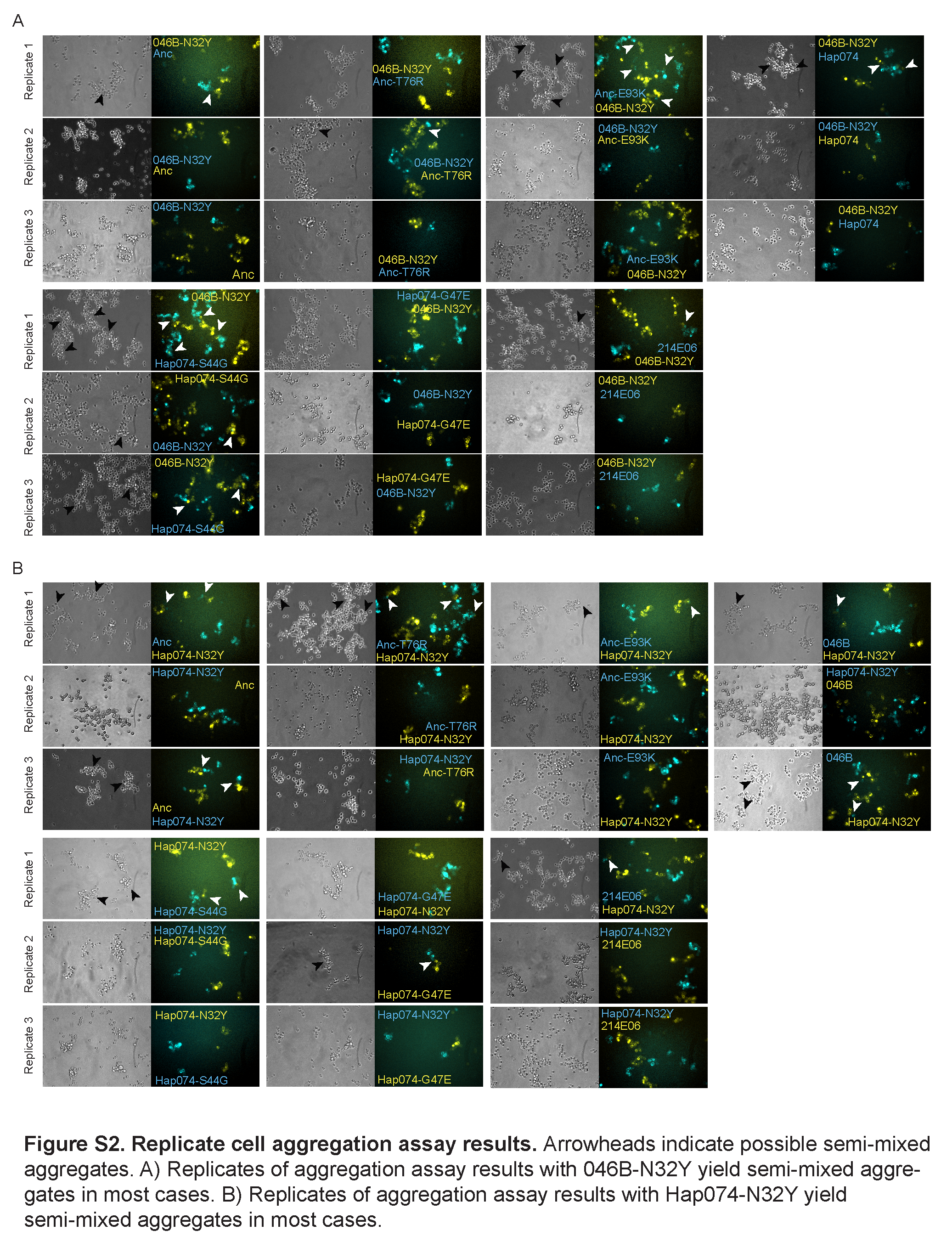

### Figure S3

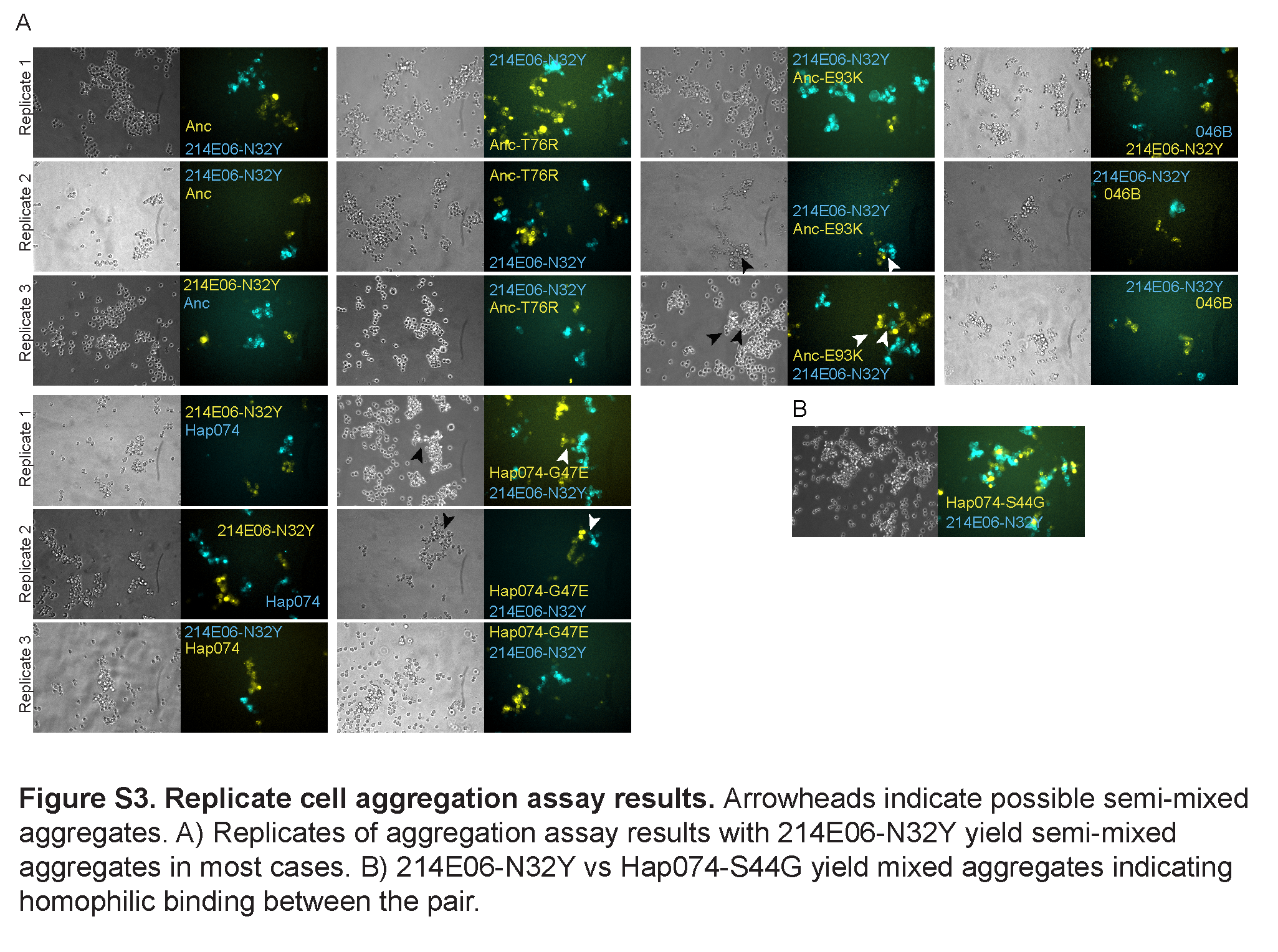
